# Rapid degeneration and neurochemical plasticity of the lateral geniculate nucleus following lesions of the primary visual cortex in marmoset monkeys

**DOI:** 10.1101/2023.09.13.557539

**Authors:** Gaoyuan Ma, Jonathan M. Chan, Katrina H. Worthy, Marcello G.P. Rosa, Nafiseh Atapour

## Abstract

Lesions of the primary visual cortex (V1) cause retrograde neuronal degeneration, volume loss and neurochemical changes in the lateral geniculate nucleus (LGN). Here we characterise the timing of these processes, by comparing the LGN of adult marmoset monkeys following various recovery times after unilateral V1 lesions. Observations in NeuN-stained sections obtained from animals with very short (3 days) recoveries showed that the volume and neuronal density in the LGN ipsilateral to the lesions were similar to those in the contralateral hemispheres. However, neuronal density in the LGN lesion projection zones (LPZ) dropped rapidly thereafter, with a loss of ∼50% of the population occurring within a month of the lesions. This level of neuronal loss was then sustained for the remainder of the range of recovery times, up to >3 years post-lesion. In comparison, shrinkage of the LGN progressed more gradually, not reaching a stable value until 6 months post lesion. We also determined the time course of the expression of the calcium-binding protein calbindin (CB) in magnocellular (M) and parvocellular (P) layer neurons, a recently described form of neurochemical plasticity triggered by V1 lesions. We found that CB expression could be detected in surviving M and P neurons as early as 1 month after a lesion, with the percentage of neurons showing this neurochemical phenotype showing subtle changes over 6 months. Our study shows that there is a limited time window for any possible interventions aimed at reducing secondary neuronal loss in the visual afferent pathways following damage to V1.

## Introduction

Lesions in the primary visual cortex (V1) trigger degeneration and volume loss in the lateral geniculate nucleus (LGN), both in human and non-human primates (Atapour et al., 2017, 2021; Cowey & Stoerig, 1989; Hendrickson et al., 2015; Simmen et al., 2022). The widespread degeneration involves projection neurons in the magno-(M), parvo (P), and koniocellular (K) neurons as well as corticogeniculate fibers (Atapour et al., 2017; Kinoshita et al., 2019; Wong-Riley, 1972). V1 lesions also cause cortical blindness, a condition that may reflect not only the immediate loss of cortical tissue, but also the secondary loss of LGN and retinal cells (Leopold, 2012).

However, despite degeneration and volume loss caused by the lesion, there are a number of surviving neurons in the LGN, offering the potential for plasticity and recovery (Yu et al., 2018). One recently revealed form of plasticity is changes in protein expression in the surviving neurons (Atapour et al., 2021, 2022). Neurons in the normal LGN show specificity in their expression of calcium-binding proteins, with M and P neurons exclusively expressing parvalbumin (PV) and K neurons expressing calbindin-D28K (CB; Yan et al., 1996). In animals with long-term V1 lesions, this specificity is disrupted with many M and P neurons co-expressing CB and PV (Atapour et al., 2022). It has been found that some of the neurons undergoing this neurochemical change form projections to the middle temporal area (area MT) (Atapour et al., 2022), one of the key cortical areas hypothesized to mediate residual visual capacities following V1 lesions (Ajina & Bridge, 2018; Rosa et al., 2000).

Retrograde degeneration following V1 lesions is likely to be a gradual process (Atapour et al., 2017; Mihailović et al., 1971; Wong-Riley, 1972, Cowey et al., 1999, 2011). However, data on the exact pace of the degeneration remains sparse, particularly in the first weeks after V1 lesions. In addition, little is known about the time course of changes in calcium-binding protein expression. Here we address this knowledge gap by studying the cellular anatomy of the LGN in marmoset monkeys in the first weeks or months following V1 lesions. Marmosets are non-human primates which have been gaining prominence as a translational model for studies of human biology and disease (Mitchell & Leopold, 2015; Okano, 2021; Solomon & Rosa, 2014). Understanding the rate of progress of post-injury degenerative processes may provide important clues for future work aimed at minimising cell loss, and potentially greater preservation of residual visual abilities.

## Materials and Methods

### Subjects

The present study was performed in a cohort of 20 marmoset monkeys (*Callithrix jacchus*), of which 13 received a unilateral (left hemisphere) V1 lesion in adulthood and 7 were non-lesioned controls (see Table 1). Eleven of these animals (4 lesioned, 7 controls) were part of other experiments involving fluorescent tracer injections (Majka et al., 2020) or electrophysiology experiments, not reported here. All animals were kept within family groups and lived in large cages with access to both indoor and outdoor environments. Their health condition and well-being were monitored throughout the experimental period. The experiments were conducted based on the Australian Code of Practice for the Care and Use of Animals for Scientific Purposes. All experiments were approved by the Monash University Animal Ethics Experimentation Committee.

**Table 1.** Experimental details for all subjects.

| Subject (Sex) | Survival<br>(~months) | Shrinkage ratio<br>(% ipsi/intact) | Age at<br>lesion<br>(~months) | Age at<br>perfusion<br>(~months) | Analysed<br>LGN |
| --- | --- | --- | --- | --- | --- |
| WA9(M) | 39 | 50.84% | 19 | 58 | Both |
| WA14*(F) | 31 | 52.30% | 25 | 57 | Both |
| WA13*(M) | 28 | 68.61% | 29 | 57 | Both |
| WA16<sup>#</sup>(M) | 23 | 57.36% | 32 | 55 | Both |
| WA15*(F) | 12 | 43.17% | 26 | 38 | Both |
| WA25(F) | 6 | 71.53% | 26 | 32 | Both |
| WA22(F) | 3 | 85.11% | 42 | 45 | Both |
| WA23(F) | 3 | 85.98% | 37 | 40 | Both |
| WA20(M) | 2 | 75.87% | 36 | 38 | Both |
| WA21(F) | 2 | 84.02% | 44 | 46 | Both |
| WA27(M) | 1 | 78.83% | 39 | 40 | Both |
| WA24(F) | 3 days | 96.65% | 32 | 32 | Both |
| WA26(M) | 2 days | 101.58% | 45 | 45 | Both |
| CJ227*(M) | - | 102.51% | - | 37 | Both |
| CJ217*(M) | - | 99.37% | - | 34 | Both |
| CJ200*(F) | - | 98.39% | - | 44 | Both |
| F1741*(F) | - | - | - | 42 | Right |
| CJ174<sup>#</sup>(F) | - | - | - | 32 | Right |
| CJ170<sup>#</sup>(M) | - | - | - | 29 | Right |
| CJ167<sup>#</sup>(F) | - | - | - | 28 | Right |

*\* Animals that received fluorescent tracer injections for other projects. # Animals that participated in electrophysiology experiments for other projects. CJ217 and CJ227 were only used for volume analysis. (M) Male; (F) Female.*

### V1 lesions

The V1 lesion surgery was conducted following an updated procedure based on the technique introduced by Rosa et al., (2000). This procedure involves an occipital lobectomy along a vertical plane along the border between V1 and the second visual area (V2; Rosa et al. 1997), resulting in a complete loss of the representation of the visual field up to 10° eccentricity along the vertical meridian, and 20-30° along the horizontal meridian.

Reconstructions of lesions and visual field defects created with this procedure can be found in earlier publications from our group (e.g. Yu et al. 2013, 2018; Atapour et al. 2021).

The animals were pre-medicated with oral meloxicam (Metacam; Boehringer Ingelheim, 0.1 mg/kg, i.m) and cephalexin (Ibilex; Alphapharm P/L, 30 mg/kg, i.m) 24 hours before the surgery. Atropine (Atrosite; Ilium, 0.2 mg/kg) was administered 30 minutes before anaesthesia, which was accomplished by inhalation of isoflurane (Isorrane; 4 to 5% in oxygen, Baxter). Dexamethasone (Dexason; Ilium, 0.3 mg/kg, i.m) was also administered. During the surgery, the animals were positioned in a modified stereotaxic head holder while their heart rates, body temperatures, and body oxygenation levels (PO_2_) were continually monitored. The anaesthetic condition was adjusted continuously (isoflurane, between 2 to 5%) to ensure the animals showed no spontaneous muscle activity and had no withdrawal reflexes. Following a left hemisphere craniotomy and durectomy the occipital pole was removed using a fine-tipped cautery, following a vertical excision along a plane extending from the dorsal surface of the occipital lobe to the cerebellar tentorium, across its entire mediolateral extent. After the removal of tissue, the resulting cavity was filled with haemostatic microspheres (Arista AH, BARD Davol Inc.) until the bleeding stopped. The surface of the wound was covered with ophthalmic film (Gelfilm, Pfizer Inc.), and the cavity was filled with Gelfoam (Pfizer Inc.). The skull flap was repositioned and secured with cyanoacrylate (Vetbond, 3M), followed by skin suture with polyglactin thread (5-0 Vicrly, Johnson & Johnson). The animals then were placed in an infant incubator (Atom Medical) for recovery and reintroduced to the home cage after recovery of mobility. During recovery, postoperative analgesia (oral meloxicam 0.05 mg/kg for adults, 3 d) and antibiotic (oral cephalexin 30 mg/kg, 5 d) was given. The animals demonstrated normal movement abilities including precise grasping, holding, jumping between branches, and obtaining food without assistance already on the day following surgery, and throughout the recovery period.

### Tissue processing

Following variable survival periods (Table 1) the animals were anaesthetised with Alfaxan (Ibilex, 30mg/kg i.m.) and then overdosed with a pentobarbitone sodium injection (100 mg/kg i.v.). Following cardiac arrest, they were perfused with 0.1 M heparinised phosphate buffer (PB; pH 7.2) and 4% paraformaldehyde (PFA) in 0.1 M PB. The brain was removed and post-fixed for 24 hours in the same solution, after which cryoprotection was performed by immersion in 4% buffered PFA solutions with increasing concentrations of sucrose (10%, 20% and 30%). The brain was then snap-frozen and cut into 40μm coronal sections using a cryostat (Leica, CM1850).

For NeuN staining, every fifth section was washed in a PB solution (0.1M). The sections were incubated with a blocking solution (0.3% Triton-X100 and 10% horse serum in 0.1 PB solution) for 1 h at room temperature, and then treated with NeuN primary antibody (1:800, Merck Millipore, USA, MAV377, Clone A60, RRID: AB_2298772) for 46-48 h at 4°C. This was followed by incubation with biotinylated horse anti-mouse IgG secondary antibody (1:200, PK-6102, Vectastain Elite ABC HRP kit, Vector Laboratories, USA, RRID: AB_2336821) for 30 min at room temperature. The sections were then incubated with ABC reagent (100µl of solution A and 100µl of solution B in 5 ml of 0.1M PBS) for 30 min at room temperature. After incubation, sections were stained with DAB substrate working solution (DAB kit SK-4100, RRID: AB_2336382) for 30 min at room temperature, followed by three washes with PB (0.1M). The mounted sections were dried for 48 h before being dehydrated in ethanol and xylene, and coverslipped with DPX.

For fluorescence staining of calcium-binding proteins another 1 in 5 series of sections was incubated with blocking solution for 1 h at room temperature, followed by 46-48 h incubation in primary antibodies for CB (1:50, Thermo Fisher, USA, RRID: AB_2068509) and PV (1:1000, Swant, Switzerland, code 235, RRID: AB_10000343). The secondary antibodies (Alexa Flour 488, 1:600, ab150109, Thermo Fisher, or Alexa Fluor 647, 1:600, ab150064, Thermo Fisher) were applied for 60 min at room temperature, followed by coverslip using antifade mounting medium (Vectashield, Vector Laboratories).

### Quantification and statistical analysis

For NeuN-stained brain sections, slides were scanned with an Aperio Scanscope AT Turbo microscope (Leica Biosystems) at ×20 magnification under the resolution of 0.50 μm/pixel, and analysed using Aperio Image Scope software. Estimation of volume was done using Cavalieri estimator based on the area of LGN in equally spaced sections throughout the anteroposterior extent of this nucleus (Atapour et al., 2017; Gundersen & Jensen, 1987). For neuronal density, counting frames (150×100μm^2^) were placed systematically on each LGN section covering both degenerated and undegenerated sectors. In each section, 2-12 of counting frames were placed in a radial direction across all layers (Figure 1). Only neurons with clear staining were counted, regardless of shape. Every neuron that was fully located inside the frame or that intersected the top or right edges was included. Data of all sections were combined for analysis. The cell counts were converted to densities (cells/ mm^3^) by taking into consideration the section thickness and a shrinkage factor of 0.801 (Atapour et al., 2019).

**Figure 1.**
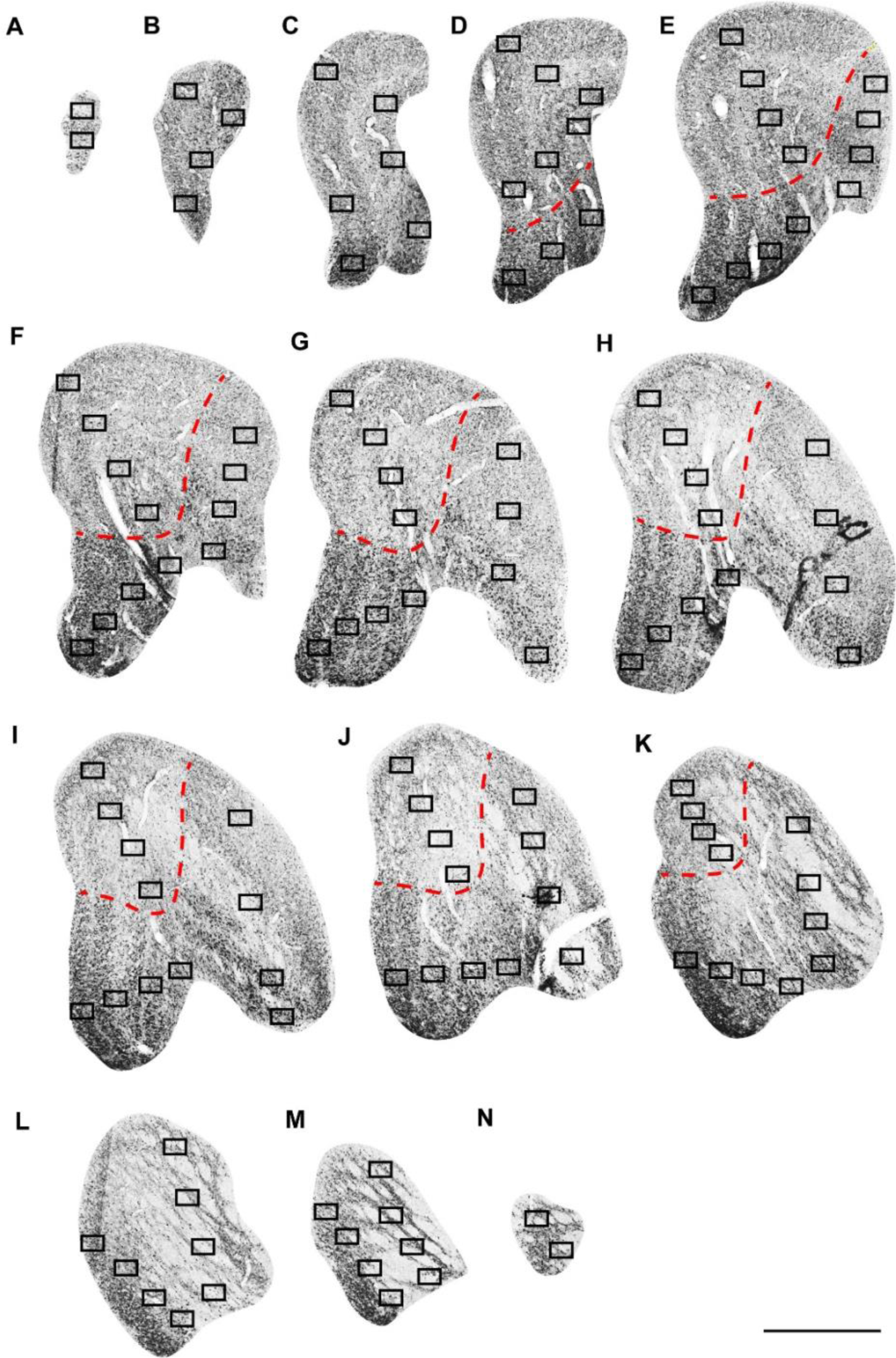
Design of sampling for stereological analysis. **A-N:** Representative coronal sections covering the extent of lateral geniculate nucleus (LGN, interaural +3.15mm to +6.00mm, Paxinos et al., 2012), on which location and number of sampling frames (100 × 150 µm) are shown. **A** represents the most posterior part of LGN. Red dashed lines separate the lesion projection zones (LPZ) from the rest of LGN. The counting frames are placed in a radial dimension covering the different cellular layers in LGN. Scale bar: 1mm.

The fluorescence immunostained sections were scanned using confocal microscopes (Nikon C1 invert, 20x magnification, filters 488, 647) and analysed with ImageJ software (Fiji, USA). The LGN was sampled in fixed sized square frames (638 x 638μm^2^) from the middle sections of the LGN, where the placement of the counting frames covered most of LGN as introduced in our previous study (Atapour et al., 2021). Only neurons with clear fluorescent signal were counted using the cell counter plug-in. The analysis was centred on determining the percentages of neurons expressing CB, PV, or both calcium-binding proteins

All statistical analysis was performed using Prism 9 (GraphPad software, USA). Data were analysed using student’s t-test, one- or two-way ANOVA and linear regression where applicable and presented as mean ± standard error of the mean (SEM). Statistical results with a p value < 0.05 were considered statistically significant.

## Results

### V1 lesion triggers neuronal loss in the LGN within a month

In the animals perfused 2-3 days post lesion NeuN staining of the LGN appeared normal in terms of volume, lamination and neuronal density, with no differences being observed between the LGNs ipsilateral and contralateral to the lesions (Figure 2). In all other cases (e.g. Figure 3 a pale staining region corresponding to the lesion projection zone (LPZ) was obvious in the ipsilesional LGN, similar to previous observations (Atapour et al., 2017; Hendrickson et al., 2015; Kisvárday et al., 1991). Higher magnification views of neurons within and outside the LPZs are shown in Figure 4A, showing that the pale staining regions were characterised by obvious reduction in neuronal density.

**Figure 2.**
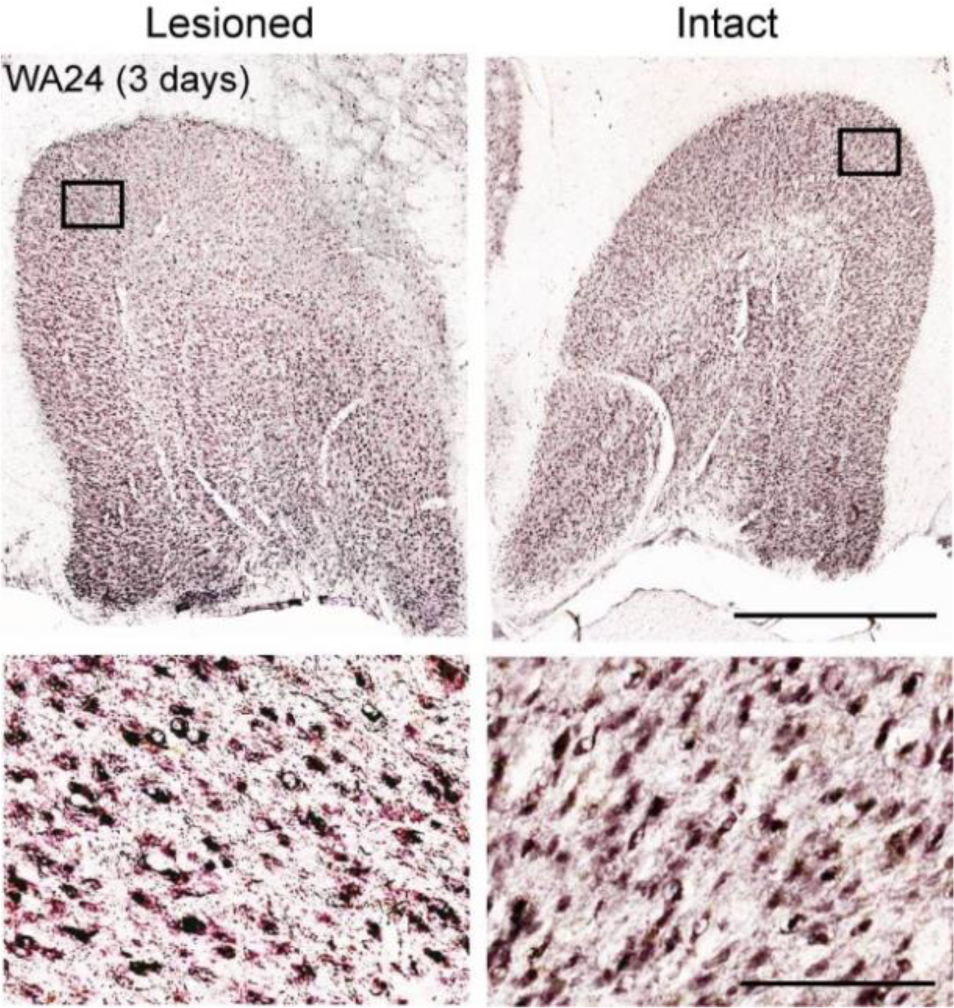
No cellular loss is observed in the lateral geniculate nucleus (LGN) three days after lesion in the primary visual cortex (V1). **Top:** LGN in the lesioned (Left) or intact hemispheres (right), Scale 1 mm. **Bottom:** Boxed areas in the expected lesion projection zone (parvocellular layers) and a corresponding region of the contralateral hemisphere are shown in higher magnification. Scale bar= 250μm.

**Figure 3.**
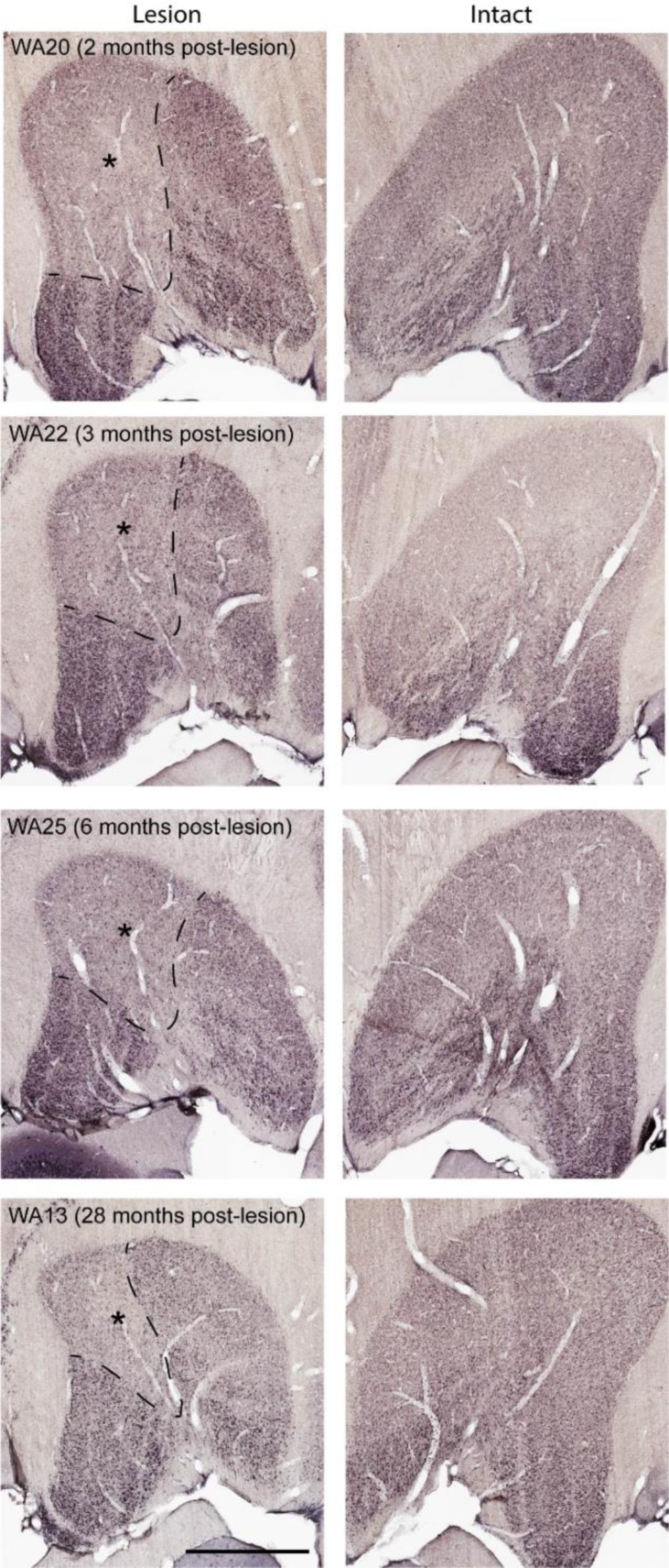
Lesions of the primary visual cortex (V1) trigger retrograde degeneration in the lateral geniculate nucleus (LGN). **A:** Representative images of NeuN-stained coronal sections through the LGN (interaural +4.80mm), obtained from four animals with different post-lesion survival times. In all cases the LGN ipsilateral to the V1 lesion shows a well-defined lesion projection zone (LPZ, identified by dashed line/ asterisk), while the contralateral LGN shows normal lamination and shape. Scale bar: 1mm.

**Figure 4.**
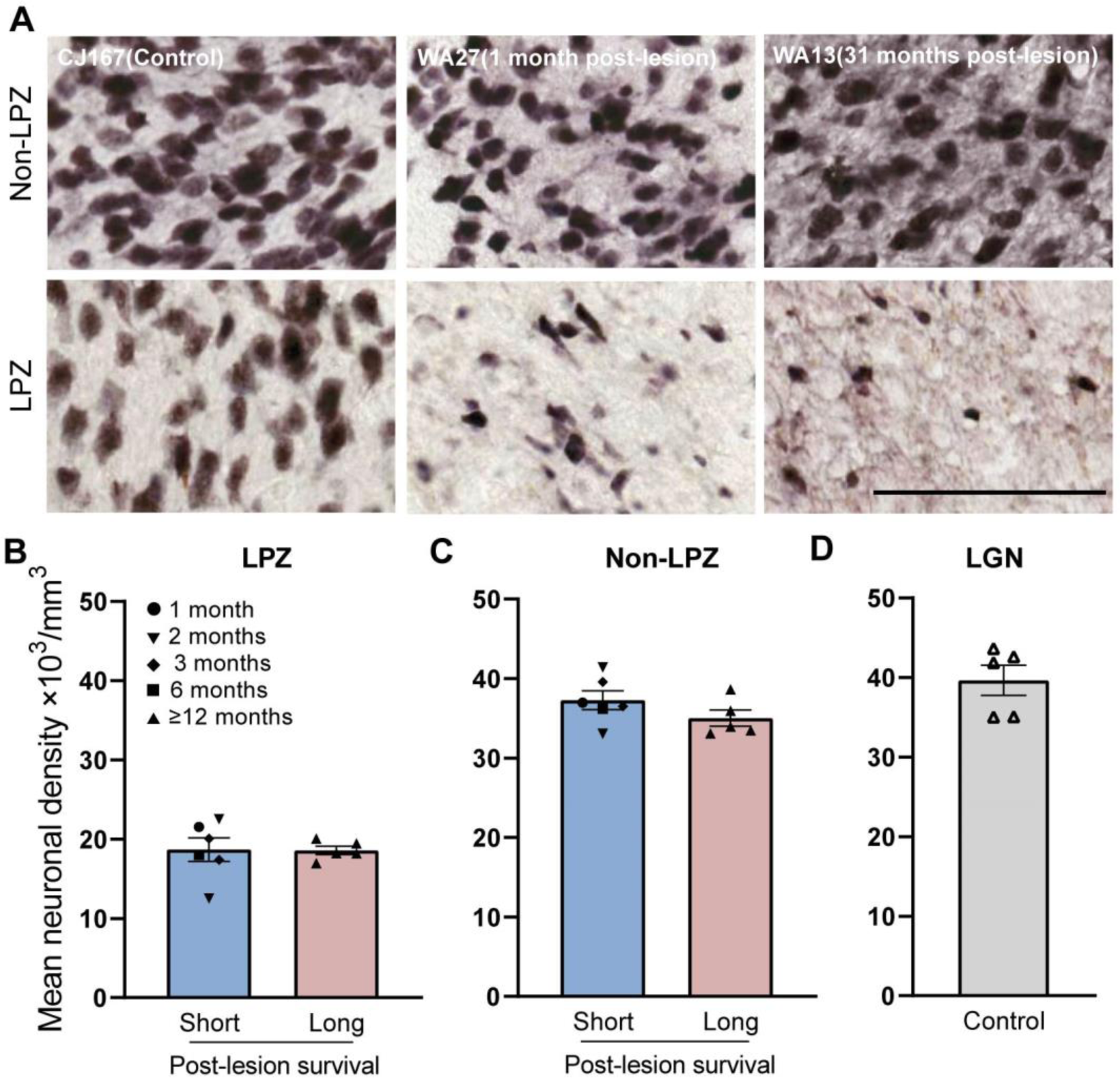
Lesions of the primary visual cortex (V1) cause significant neuronal loss in the lateral geniculate nucleus (LGN) within the first month. **A:** Representative images from the lesion projection zones (LPZs) and non-LPZ regions in control (CJ167), short survival (WA27) and long survival (WA13) cases. Scale bar = 100 µm. **B-D:** Mean ± SEM of neuronal density of LGN in the short and long survival animals for LPZ (B), non-LPZ (C) and control (D) sectors. Different post-lesion survival times are identified with different symbols in the lesioned cases.

For analysis, the cohort of animals was divided into short survival (<6 months) and long survival (12-39 months) groups, the latter corresponding to time points already explored in our earlier studies (Atapour et al., 2022; Chan et al., 2021). As shown in Figure 4B, the neuronal density in the LPZs dropped to almost half of the values observed outside the LPZs and control animals. This drop was evident even after a month post lesion (Case WA27), indicating a rapid neuronal loss following V1 lesion. In fact, the observed neuronal loss was similar among all cases in the short survival group (survivals between 1 and 6 months (Table 1, Figure 4B). Comparison with those in the long survival group also did not reveal any differences for neuronal density in the LPZ, indicating no further degeneration (Figure 4B, C [two-way ANOVA; survival time: *F* (1, 18) =1.02, p=0.33, zone: *F* (1, 18) =225.1, p< 0.0001, interaction; *F* (1, 18) =0.86, p=0.37, post hoc; LPZ vs. non-LPZ; short survival (18.68±1.47 vs 37.28±1.18 ×10^3^/mm^3^, p<0.0001), long survival (18.59±0.54 vs. 35.02±1.03 ×10^3^/mm^3^, p<0.0001)]. The mean neuronal density outside the LPZ was similar to that found in the LGN of non-lesioned control animals, both in the short and long survival groups (Figure 4C, D, One-way ANOVA, F (2, 13) = 2.582, p=0.11).

### LGN shrinkage advanced with the time after V1 lesion

The average LGN volume loss (ratio of ipsilesional LGN volume to contralesional LGN volume) was more extensive in th4 long survival group compared to the short survival group (Figure 5A, t-test; long survival vs. short survival, 54.46±4.21 vs. 80.22±2.36%, p = 0.0003). To account for the possibility variations in lesion sizes, we ran a separate analysis comparing the percentage loss of volume in the LPZ and non-LPZ regions, compared to the contralesional LGN volume. This analysis showed that the difference between the two groups originates mainly from changes in the LPZ volume (Figure 5B; long vs. short survival, LPZ; 17.93±2.02 vs. 30.85±1.41%, p = 0.0004). Although the analysis of non-LPZ volume shown in Figure 5C indicates some variability across cases, the results did not reach statistical significance (non-LPZ; 36.52±6.12 vs. 47.26±3.18%, p = 0.13). In summary, the result indicates a significantly smaller volume loss in the short survival group.

**Figure 5.**
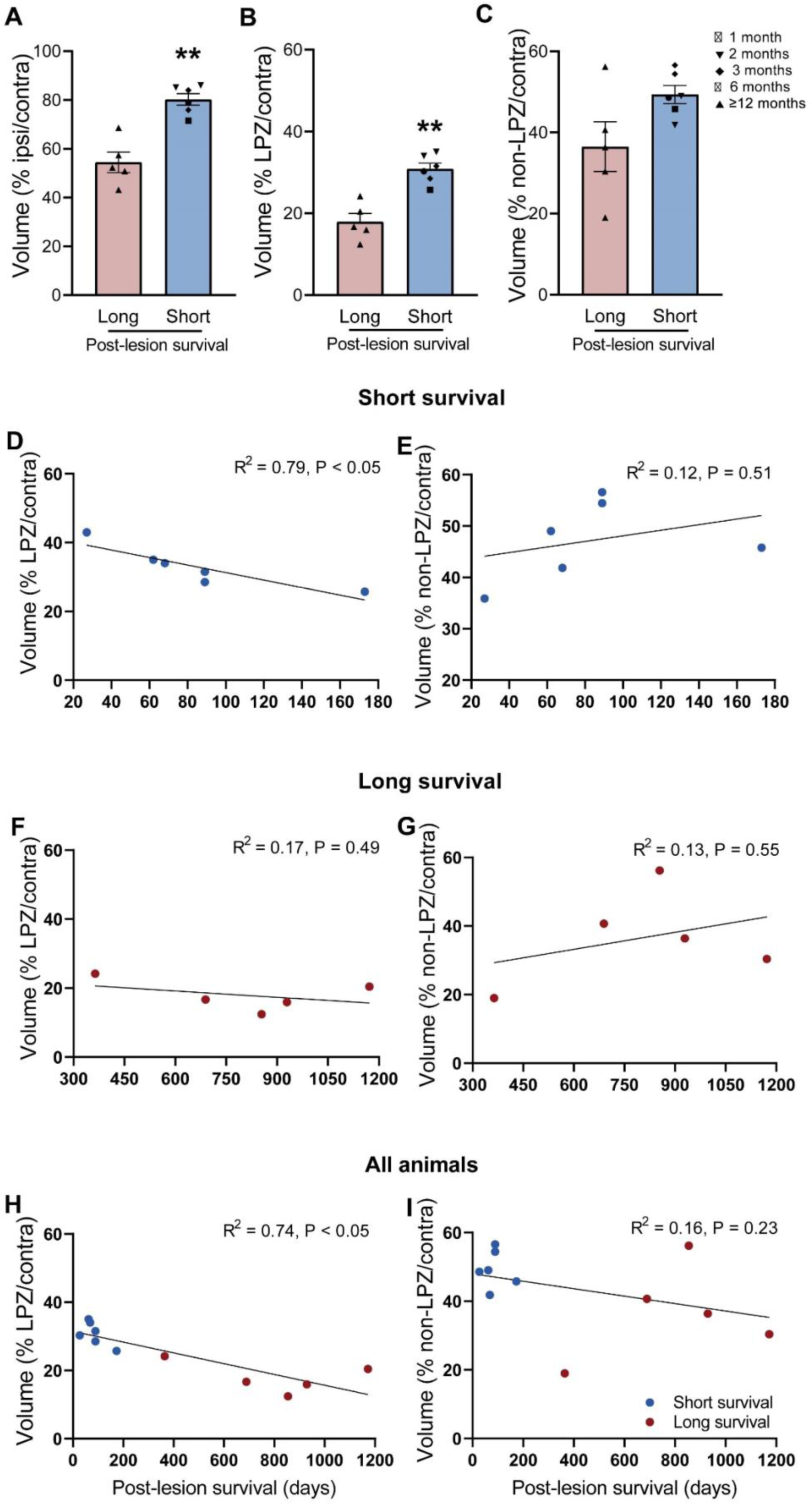
Lateral geniculate nucleus (LGN) volume loss correlates with the post-lesion survival time. **A-C:** Mean ± SEM of volume of the LGN for animals with short (1-6 months, n = 6) and long (12-39 months, n = 5) post-lesion survival period. Different post-lesion survival times are identified with variable symbols in the short survival cases. ** p < 0.01. **D-I**: Percentage mean volume of lesion projection zone (LPZ, D, F, H) or non-LPZ (E, G, I) sectors are plotted against the post-lesion survival time. Data obtained in the short survival and long survival groups are shown separately (D-G) or combined (H-I). R^2^ represents the significance levels for the slopes of simple linear regressions for each condition.

To further understand the differences in volume loss in relation to variable survival times, we plotted percentage volume (i.e. the ratio of the volumes in the ipsilesional vs. contralesional LGN) against the survival time post-lesion in Fig 5 (D-I). A negative linear correlation was observed for LPZ volume (compared to contralesional LGN, Fig 5D, linear correlation, R^2^ = 0.73, p=0.0016). When the correlations were conducted separately for the short and long survival group, only the short survival group demonstrated a statistically significant negative correlation (linear correlation, LPZ; Fig 5D; R^2^ = 0.79 p=0.04, 4F; R^2^ = 0.17 p = 0.49). This finding is consistent with our previous study that the shrinkage of LGN stabilised within six to seven months following V1 lesions (Atapour et al., 2017). The normalised volume of non-LPZ (as compared to contra-lesion LGN), on the other hand, did not show statistically significant changes in both short and long survival groups (linear correlation, non-LPZ; Fig 5G; R^2^ = 0.14 p=0.29, 5H; R^2^ = 0.02 p = 0.83, 5I; R^2^ = 0.13 p = 0.55). Thus, the results indicate a substantial rapid loss of neurons occurs first, followed by a gradual shrinkage of the LGN after lesion, and argue against differences in lesion extent being a confounding factor.

### Calbindin positive M and P neurons exist as early as one-month post lesion

Our previous studies demonstrated that, in addition to degeneration, V1 lesions elicit neurochemical changes in the LGN M and P neurons (Atapour et al., 2022). In particular, these neurons, which normally only express PV, also show co-expression of CB, including in M neurons projecting to area MT (Atapour et al., 2022). Here we explored the timing of CB expression in M and P neurons following V1 lesions. The CB positive neurons were quantified from populations of PV-expressing (M and P neurons) in the short survival group. This confirmed the presence of CB in all cases, ranging from 1 to 6 months post-lesion (Figure 6). Example images of CB expression in M and P neurons are shown in Fig 6A. While degeneration is exclusive to the ipsilesional LGN, a minimum of 11% CB co-localisation with PV was also detected in the contralateral LGN (Figure 6B), confirming that this neurochemical change triggered by the unilateral lesion occurs in both hemispheres (Atapour et al, 2022). The average values obtained from all animals showed that the proportion of CB expressing M and P neurons was significantly higher in the ipsilesional LGN (Fig 6B, paired t-test; ipsi vs contra; 23.53±2.62 vs. 19.37±2.98). Regression analysis revealed a trend towards greater CB expression in M and P neurons with the survival time, although this did not reach a statistically significant level (Figure 6C, R^2^ = 0.60 p = 0.07).

**Figure 6.**
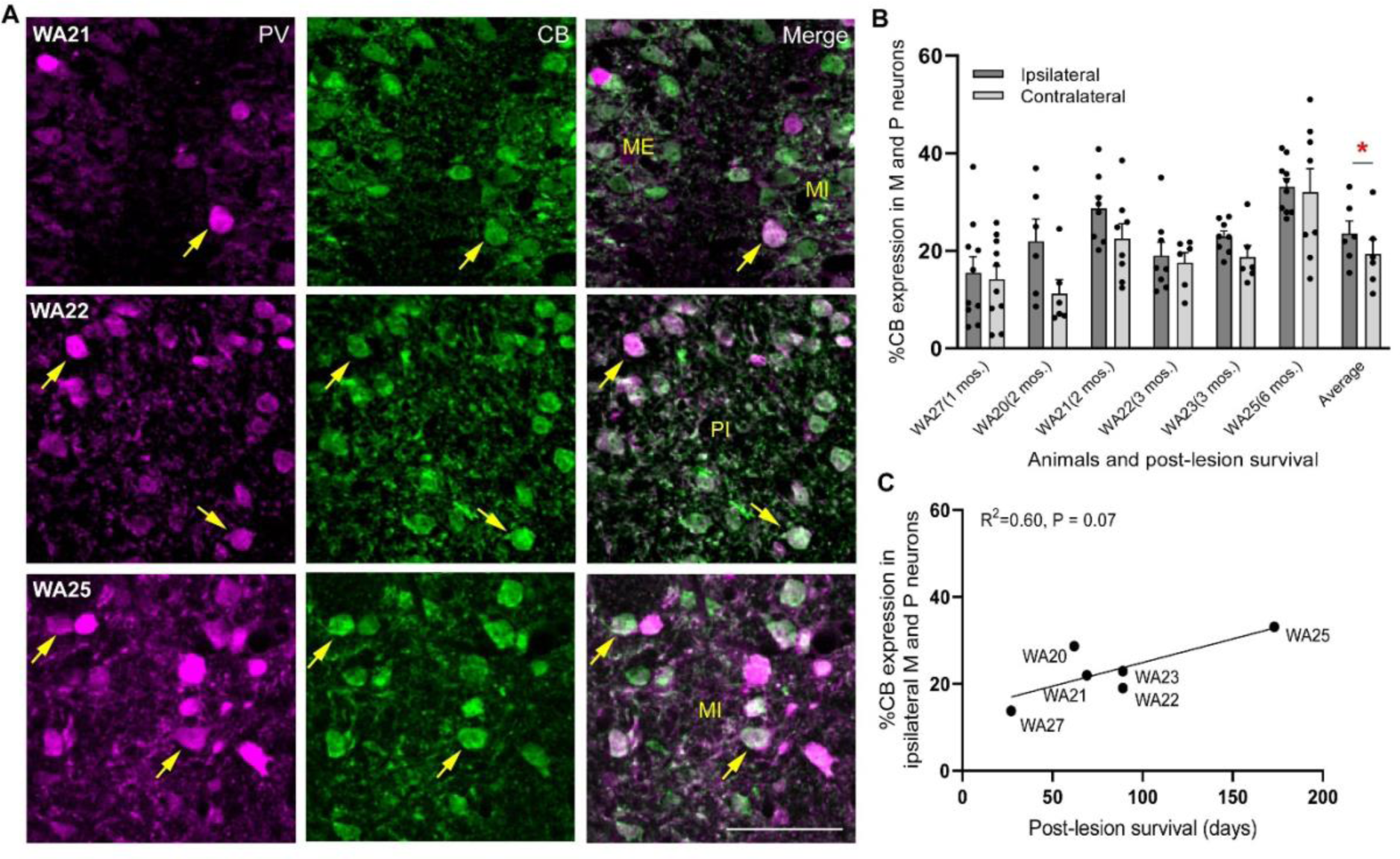
Calbindin (CB) immunoreactivity in parvalbumin (PV)-expressing neurons of lateral geniculate nucleus (LGN) is present after a month following lesions of the primary visual cortex (V1). **A:** Representative images showing CB expression in PV-expressing magnocellular (M) and parvocellular (P) neurons for 3 animals with two (WA21), three (WA22) and six (WA25) months post-lesion survival. Yellow arrows point to individual neurons expressing both CB and PV. ME: magnocellular external layer, MI: magnocellular internal layer, PI: parvocellular internal layer. Scale bar: 100μm. **B:** Percentage mean ± SEM of CB expression in M and P neurons of the LGN (identified by PV staining) within six months post lesion. Black circles represent individual sections analysed. **C:** Mean percentage of CB expression in the ipsilesional M and P neurons is plotted against post-lesion survival time. R^2^ indicates the significance level for the slope of a simple linear regression.

## Discussion

The present study quantified changes in the structure and neurochemistry of the primate LGN within the first six months following V1 lesions. We found that the vast majority of the changes in LPZ neuronal density occurred within a month of the lesion, with no further changes occurring up to 3 years post lesion. However, shrinkage of the LPZ happened in a slower timescale, being completed within 6 months post lesion. In parallel to these degenerative changes, we found that CB expression could be detected in surviving M and P neurons as early as one-month post lesion.

### Neuronal loss in the LGN

V1 lesion induced fast retrograde degeneration, which resulted in a sharp decrease of neuronal density in the LGN just one-month post lesion. The fast nature of degeneration is in agreement with previous studies in other primate species (Mihailović et al., 1971; Wong-Riley, 1972), which used the Nissl stain to assess neuronal loss. In squirrel monkeys, neuronal degeneration is reported to be completed within 42 days, with signs of retrograde degeneration being already evident only 5-8 days after lesions (Wong-Riley, 1972). This time course is similar to that suggested by our data, where no sign of cell loss was detected 3 days after lesion. In macaque monkeys, previous work indicating that the process is completed within 3– 6 weeks (Mihailović et al., 1971).

In the present study the small variability in the estimates of neuronal density in the LPZ among cases in the short survival group, together with the similar values obtained in the long survival group, indicate that the degeneration process has been largely completed by the end of the first month post lesion. In the context of previous literature, these results are significant in two ways. First, they demonstrate that the time course of degeneration is similar despite differences in brain size (and hence axonal length) among primates. This species similarity indicates that a comparable time course is likely to apply to the human brain. Second, they identify a time window during which possible interventions to reduce neuronal loss in the LGN need to apply. Long-term occipital lesions cause loss of function not only due to the direct loss of V1 cortex, but also subsequent degenerative changes in the LGN and retina (Atapour et al., 2017, 2021; Cowey et al., 2011; Hendrickson et al., 2015). Preventing or minimising the secondary losses in the afferent visual pathway may be relevant for future therapeutic interventions aimed at retaining or improving residual vision.

Atrophy of the LGN is a critical pathological indication for V1 lesions which can be quantified with non-invasive brain imaging (Simmen et al., 2022). Whereas volume reduction is clearly influenced by neuronal loss, these two aspects of degeneration can occur independently. For example, volume changes without concomitant loss of neuronal cell bodies have been observed in the pulvinar complex following V1 lesions, an observation that has been interpreted as indicating that volume reduction was likely due to changes in the neuropil (Chan et al. 2021).

In the present study, the volume loss in the LGN ipsilateral to the lesion increased with the survival time, despite the level of neuronal loss being similar across all cases. This observation was normalised for the lesion size (a factor which is likely to affect LGN shrinkage) by separate estimation of percentage volume loss for LPZ and non-LPZ. This correlation with survival time was not observed among long survival animals (Fig 4). This is consistent with previous observations (Atapour et al., 2017) that volume loss in the marmoset LGN does not progress beyond 6-7 months post lesion.

The delayed time scale for volume loss, compared to neuronal degeneration, may be due to the time required for debris removal (Liu et al., 2022; Westman et al., 2019), neuropil loss, or even remodelling of the capillary bed due to reduced energy requirements following apoptosis of LGN neurons. Moreover, cortico-geniculate axons and terminals fill much of the space between LGN neurons (Briggs & Usrey, 2011; Ichida et al., 2014), and their loss (Briggs & Usrey, 2011; Ichida et al., 2014) may also contribute to volume changes. Although evidence suggest that the time course and magnitude of retrograde and Wallerian (anterograde) degeneration are similar (Kanamori et al., 2012), there may be differences due to the differential site of injury along the axons (Lubińska, 1982) or heterogeneity of axons and their compartmentalisation (Conforti et al., 2007). Although it has been reported that early signs of cortico-geniculate terminal degeneration precede the retrograde degeneration of LGN neurons in squirrel monkeys (Wong-Riley, 1972), the biological complexity of degenerative processes (Conforti et al., 2007) may create difficulty in pinpointing the precise timing of events. What is clear is that the physical collapse of LGN volume is a more prolonged process than neuronal death.

### Expression of CB in surviving neurons

Recent studies have shown that neurochemical changes are part of long-term plastic changes in the primate LGN following V1 injury (Atapour et al., 2022). Here, we show evidence that CB expression in M and P neurons is evident even a month after lesion. However, the percentage of M and P neurons expressing CB is lower (15-35%) than that observed in longer survival times (35-75%, Atapour et al., 2022), reiterating the gradual nature of the emergence of CB expression in M and P neurons. One particularly puzzling aspect of this process is the fact that changes in the neurochemistry of M and P neurons are bilateral, which may point to more global changes in the excitation-inhibition network (Atapour et al. 2022). CB expression has been highlighted as a protective mechanism linked to neuronal resilience under stress (Ng et al., 1996). Thus, concomitant with the ongoing degeneration in LPZ, the neurons in non-LPZ undergo neurochemical changes that may prepare them for better survival, or regenerative processes (Choi et al., 2008; Kim et al., 2006; Zupanc & Zupanc, 2006). As recently demonstrated, the latter involves remodelling of the projections of M neurons, which form projections to extrastriate cortex (Atapour et al., 2022).

### Limitations and suggestions for future studies

The present study focuses on the changes occurring in LGN following V1 lesions in young adult marmosets. However, there is evidence that the age at which injury happens could affect the extent and pace of the degeneration process (Atapour et al., 2017; Dineen & Hendrickson, 1981; Hendrickson et al., 2015; Weller & Kaas, 1989). Thus, additional work focused on different stages of life is advisable in order to obtain a full picture of the effects of V1 lesions. Earlier work comparing lesions in young (2-6 weeks postnatal) and adult marmosets revealed modest physiological differences in the surviving LGN neurons (Yu et al. 2018).

The extent of the damage to the extrastriate visual cortex can also influence estimates of degeneration (Cowey et al., 1999). In the present experiments, the lesions were designed to be largely contained to V1, although involvement of parts of the second visual area near the border is inevitable. More extensive lesions resulting in damage to rostral extrastriate areas would likely result in more extensive loss due to the removal of projection targets of neurons that would otherwise survive the degeneration process. However, the few surviving neurons in the LGN continue to receive retinal projections even following complete unilateral hemispherectomies (Boire et al., 2001). Whether the surviving neurons following larger lesions undergo biochemical changes remains an open question.

## Conclusions

Our study demonstrates that the neuronal loss in the LGN proceeds rapidly in the marmoset, being largely completed within one month of V1 lesions. At the same time, it indicates that the volume loss and biochemical changes in the LGN are slower processes, highlighting the complexity of the processes triggered by such lesions. This information has broader implications for our understanding of traumatic brain injury and future efforts towards minimizing visual loss, and promoting regenerative processes which may improve the quality of residual vision in primates, including humans.

## Notes

Competing Interest Statement: The authors declare there is no conflict of interest.

### Competing Interest Statement

The authors have declared no competing interest.

## Bibliography

Ajina, S., & Bridge, H. (2018). Blindsight relies on a functional connection between hMT+ and the lateral geniculate nucleus, not the pulvinar. PLoS Biology, 16(7), e2005769. 10.1371/journal.pbio.2005769

Atapour, N., Worthy, K. H., Lui, L. L., Yu, H.-H., & Rosa, M. G. P. (2017). Neuronal degeneration in the dorsal lateral geniculate nucleus following lesions of primary visual cortex: Comparison of young adult and geriatric marmoset monkeys. Brain Structure and Function, 222(7), 3283–3293. 10.1007/s00429-017-1404-4

Atapour, N., Worthy, K. H., & Rosa, M. G. P. (2021). Neurochemical changes in the primate lateral geniculate nucleus following lesions of striate cortex in infancy and adulthood: Implications for residual vision and blindsight. Brain Structure & Function, 226(9), 2763–2775. 10.1007/s00429-021-02257-0

Atapour, N., Worthy, K. H., & Rosa, M. G. P. (2022). Remodeling of lateral geniculate nucleus projections to extrastriate area MT following long-term lesions of striate cortex. Proceedings of the National Academy of Sciences of the United States of America, 119(4), e2117137119. 10.1073/pnas.2117137119

Boire, D., Théoret, H., & Ptito, M. (2001). Chapter 24 Visual pathways following cerebral hemispherectomy. In Progress in Brain Research (Vol. 134, pp. 379–397). Elsevier. 10.1016/S0079-6123(01)34025-6

Briggs, F., & Usrey, W. M. (2011). Corticogeniculate feedback and visual processing in the primate. The Journal of Physiology, 589(Pt 1), 33–40. 10.1113/jphysiol.2010.193599

Chan, J. M., Worthy, K. H., Rosa, M. G. P., Reser, D. H., & Atapour, N. (2021). Volume reduction without neuronal loss in the primate pulvinar complex following striate cortex lesions. Brain Structure and Function, 226(7), 2417–2430. 10.1007/s00429-021-02345-1

Choi, W.-S., Lee, E., Lim, J., & Oh, Y. J. (2008). Calbindin-D28K prevents drug-induced dopaminergic neuronal death by inhibiting caspase and calpain activity. Biochemical and Biophysical Research Communications, 371(1), 127–131. 10.1016/j.bbrc.2008.04.020

Conforti, L., Adalbert, R., & Coleman, M. P. (2007). Neuronal death: Where does the end begin? Trends in Neurosciences, 30(4), 159–166. 10.1016/j.tins.2007.02.004

Cowey, A., Alexander, I., & Stoerig, P. (2011). Transneuronal retrograde degeneration of retinal ganglion cells and optic tract in hemianopic monkeys and humans. Brain: A Journal of Neurology, 134(Pt 7), 2149–2157. 10.1093/brain/awr125

Cowey, A., & Stoerig, P. (1989). Projection patterns of surviving neurons in the dorsal lateral geniculate nucleus following discrete lesions of striate cortex: Implications for residual vision. Experimental Brain Research, 75(3), 631–638. 10.1007/BF00249914

Cowey, A., Stoerig, P., & Williams, C. (1999). Variance in transneuronal retrograde ganglion cell degeneration in monkeys after removal of striate cortex: Effects of size of the cortical lesion. Vision Research, 39(21), 3642–3652. 10.1016/s0042-6989(99)00097-8

Dineen, J. T., & Hendrickson, A. E. (1981). Age correlated differences in the amount of retinal degeneration after striate cortex lesions in monkeys. Investigative Ophthalmology & Visual Science, 21(5), 749–752.

Gundersen, H. J., & Jensen, E. B. (1987). The efficiency of systematic sampling in stereology and its prediction. Journal of Microscopy, 147(Pt 3), 229–263. 10.1111/j.1365-2818.1987.tb02837.x

Hendrickson, A., Warner, C. E., Possin, D., Huang, J., Kwan, W. C., & Bourne, J. A. (2015). Retrograde transneuronal degeneration in the retina and lateral geniculate nucleus of the V1-lesioned marmoset monkey. Brain Structure & Function, 220(1), 351–360. 10.1007/s00429-013-0659-7

Ichida, J. M., Mavity-Hudson, J. A., & Casagrande, V. A. (2014). Distinct patterns of corticogeniculate feedback to different layers of the lateral geniculate nucleus. Eye and Brain, 6(Suppl 1), 57–73. 10.2147/EB.S64281

Kim, J. H., Lee, J.-A., Song, Y. M., Park, C.-H., Hwang, S.-J., Kim, Y.-S., Kaang, B.-K., & Son, H. (2006). Overexpression of calbindin-D28K in hippocampal progenitor cells increases neuronal differentiation and neurite outgrowth. FASEB Journal: Official Publication of the Federation of American Societies for Experimental Biology, 20(1), 109–111. 10.1096/fj.05-4826fje

Kinoshita, M., Kato, R., Isa, K., Kobayashi, K., Kobayashi, K., Onoe, H., & Isa, T. (2019). Dissecting the circuit for blindsight to reveal the critical role of pulvinar and superior colliculus. Nature Communications, 10(1), Article 1. 10.1038/s41467-018-08058-0

Kisvárday, Z. F., Cowey, A., Stoerig, P., & Somogyi, P. (1991). Direct and indirect retinal input into degenerated dorsal lateral geniculate nucleus after striate cortical removal in monkey: Implications for residual vision. Experimental Brain Research, 86(2), 271–292. 10.1007/BF00228951

Leopold, D. A. (2012). Primary visual cortex: Awareness and blindsight. Annual Review of Neuroscience, 35, 91–109. 10.1146/annurev-neuro-062111-150356

Liu, K. E., Raymond, M. H., Ravichandran, K. S., & Kucenas, S. (2022). Clearing Your Mind: Mechanisms of Debris Clearance After Cell Death During Neural Development. Annual Review of Neuroscience, 45, 177–198. 10.1146/annurev-neuro-110920-022431

Lubińska, L. (1982). Patterns of Wallerian degeneration of myelinated fibres in short and long peripheral stumps and in isolated segments of rat phrenic nerve. Interpretation of the role of axoplasmic flow of the trophic factor. Brain Research, 233(2), 227–240. 10.1016/0006-8993(82)91199-4

Majka, P., Bai, S., Bakola, S., Bednarek, S., Chan, J. M., Jermakow, N., Passarelli, L., Reser, D. H., Theodoni, P., Worthy, K. H., Wang, X.-J., Wójcik, D. K., Mitra, P. P., & Rosa, M. G. P. (2020). Open access resource for cellular-resolution analyses of corticocortical connectivity in the marmoset monkey. Nature Communications, 11(1), 1133. 10.1038/s41467-020-14858-0

Mihailović, L. T., Čupić, D., & Dekleva, N. (1971). Changes in the numbers of neurons and glial cells in the lateral geniculate nucleus of the monkey during retrograde cell degeneration. Journal of Comparative Neurology, 142(2), 223–229. 10.1002/cne.901420207

Mitchell, J. F., & Leopold, D. A. (2015). The marmoset monkey as a model for visual neuroscience. Neuroscience Research, 93, 20–46. 10.1016/j.neures.2015.01.008

Ng, M. C., Iacopino, A. M., Quintero, E. M., Marches, F., Sonsalla, P. K., Liang, C. L., Speciale, S. G., & German, D. C. (1996). The neurotoxin MPTP increases calbindin-D28k levels in mouse midbrain dopaminergic neurons. Brain Research. Molecular Brain Research, 36(2), 329–336. 10.1016/0169-328x(95)00266-u

Okano, H. (2021). Current Status of and Perspectives on the Application of Marmosets in Neurobiology. Annual Review of Neuroscience, 44, 27–48. 10.1146/annurev-neuro-030520-101844

Rodman, H. R., Sorenson, K. M., Shim, A. J., & Hexter, D. P. (2001). Calbindin immunoreactivity in the geniculo-extrastriate system of the macaque: Implications for heterogeneity in the koniocellular pathway and recovery from cortical damage. Journal of Comparative Neurology, 431(2), 168–181. 10.1002/1096-9861(20010305)431:2<168::AID-CNE1063>3.0.CO;2-N

Rosa, M. G. P., Tweedale, R., & Elston, G. N. (2000). Visual Responses of Neurons in the Middle Temporal Area of New World Monkeys after Lesions of Striate Cortex. The Journal of Neuroscience, 20(14), 5552–5563. 10.1523/JNEUROSCI.20-14-05552.2000

Simmen, C. F., Fierz, F. C., Michels, L., Aldusary, N., Landau, K., Piccirelli, M., & Traber, G. L. (2022). Lateral Geniculate Nucleus Volume Determined on MRI Correlates With Corresponding Ganglion Cell Layer Loss in Acquired Human Postgeniculate Lesions. Investigative Ophthalmology & Visual Science, 63(9), 18. 10.1167/iovs.63.9.18

Solomon, S. G., & Rosa, M. G. P. (2014). A simpler primate brain: The visual system of the marmoset monkey. Frontiers in Neural Circuits, 8, 96. 10.3389/fncir.2014.00096

Weller, R. E., & Kaas, J. H. (1989). Parameters affecting the loss of ganglion cells of the retina following ablations of striate cortex in primates. Visual Neuroscience, 3(4), 327–349. 10.1017/S0952523800005514

Westman, J., Grinstein, S., & Marques, P. E. (2019). Phagocytosis of Necrotic Debris at Sites of Injury and Inflammation. Frontiers in Immunology, 10, 3030. 10.3389/fimmu.2019.03030

Wong-Riley, M. T. T. (1972). Changes in the dorsal lateral geniculate nucleus of the squirrel monkey after unilateral ablation of the visual cortex. Journal of Comparative Neurology, 146(4), 519–547. 10.1002/cne.901460407

Yan, Y. H., Winarto, A., Mansjoer, I., & Hendrickson, A. (1996). Parvalbumin, calbindin, and calretinin mark distinct pathways during development of monkey dorsal lateral geniculate nucleus. Journal of Neurobiology, 31(2), 189–209. 10.1002/(SICI)1097-4695(199610)31:2<189::AID-NEU5>3.0.CO;2-7

Yu, H.-H., Atapour, N., Chaplin, T. A., Worthy, K. H., & Rosa, M. G. P. (2018). Robust Visual Responses and Normal Retinotopy in Primate Lateral Geniculate Nucleus following Long-term Lesions of Striate Cortex. The Journal of Neuroscience, 38(16), 3955–3970. 10.1523/JNEUROSCI.0188-18.2018

Zupanc, M. M., & Zupanc, G. K. H. (2006). Upregulation of calbindin-D28k expression during regeneration in the adult fish cerebellum. Brain Research, 1095(1), 26–34. 10.1016/j.brainres.2006.04.005

